# An optimized protocol for single-brain isolation of sex-defined pure mouse astrocyte cultures

**DOI:** 10.64898/2026.03.23.713747

**Authors:** Sanarya Al-Jaf, En-Hung Ai, Julia A. Wilson, Khaled S. Abd-Elrahman

## Abstract

**Background:** Primary astrocyte cultures derived from neonatal rodent cortices provide a controlled system for investigating astrocyte-specific mechanisms. However, mixed glial preparations frequently contain contaminating microglia and oligodendrocyte progenitor cells, and most existing protocols require pooling tissue from multiple mouse pups to obtain sufficient astrocyte yields. This approach is impractical as it obscures sex and genotype, limits investigations of sex dependent astrocyte phenotypes, and precludes studies in certain transgenic models. To address this gap, our protocol achieves a high astrocyte yield from a single neonatal mouse brain, enabling sex- and genotype-specific cultures without the need for pooling. Mechanical removal of oligodendrocyte progenitors combined with pharmacological depletion of microglia using a Colony Stimulating Factor 1 Receptor (CSF1R) inhibitor produces highly enriched astrocytes suitable for functional assays, including those focused on sex-specific biology.

**Methods:** Cortical tissue was isolated from a single mouse pup is mechanically dissociated in astrocyte media. Cell suspensions are transferred to poly-D-lysine–coated flasks in astrocyte media. After 10–15 days in culture, OPCs are mechanically removed by horizontal shaking and microglia are selectively depleted by incubating cultures with CSF1R inhibitor PLX5622 for 24, 48, 72 and 96 hours. After PLX treatment, media is replaced and enriched astrocytes were maintained or passaged for experimentation. The sex of the pups is determined by PCR performed on DNA extracted from tail biopsies.

**Results:** Immunocytochemical analysis for astrocyte and microglia markers (GFAP and Iba1, respectively) showed that 24 hours of PLX5622 treatment did not fully eliminate microglia from mixed glial cultures. Extending treatment to 48 hours effectively depleted microglia while minimizing cytotoxicity and astrocyte loss and produced a pure, high-yield, sex-specific primary astrocyte culture. PCR reliably enabled the sex identification of pups used in culture using DNA extracted from tail biopsies.

**Discussion:** This protocol provides an efficient and reproducible method for generating high-purity, sex-specific primary astrocyte cultures from a single mouse brain. It improves consistency and purity while eliminating the need to pool tissue, preserving sex and genotype and enabling studies in transgenic mouse lines of both sexes.

## 1. Introduction

Astrocytes are multifunctional glial cells that are indispensable for central nervous system (CNS) development and homeostasis(1). Beyond their classical roles in metabolic support and ion buffering, astrocytes contribute to synaptic regulation, neurovascular coupling, neurotransmitter clearance, and innate immune responses(1). Dysregulation of astrocyte signaling pathways is increasingly recognized as a critical driver of neurological disease pathology, including chronic neuroinflammation, synaptic dysfunction, and impaired cerebrovascular regulation(2). Consequently, in vitro astrocyte models remain essential for dissecting astrocyte-intrinsic mechanisms that govern CNS physiology and disease progression.

Primary astrocyte cultures derived from neonatal rodent cortices provide a high-yield, physiologically relevant system enriched in proliferative glial progenitors capable of differentiating into mature astrocytes in vitro. However, traditional mixed glial preparations inevitably contain heterogeneous populations of microglia and oligodendrocyte progenitor cells (OPCs). These contaminating cell types express distinct cytokine repertoires, surface receptors, and intracellular signaling pathways that can substantially influence astrocyte behaviour, transcriptomic profiles, and responses to pharmacological perturbation(3). For example, microglia-derived cytokines such as Interleukin-1β and Tumor Necrosis Factor-alpha can induce reactive astrocyte phenotypes, while OPCs and other oligodendrocyte lineage cells secrete factors that participate in bidirectional communication with astrocytes and can influence astrocyte function in the CNS (4–6). As a result, precise removal or depletion of non-astroglial cell types is critical for ensuring experimental specificity.

Several enrichment strategies have been developed, with mechanical shaking being a long-standing method to detach OPCs based on their weaker adherence to poly-D-lysine (PDL)-coated flasks. While effective for OPC reduction, mechanical dissociation alone does not adequately remove microglia, which remain adherent and functionally active within the culture (7). Advances in microglial biology have identified the colony-stimulating factor 1 receptor (CSF1R) as an essential signaling pathway for microglial survival(8). Small-molecule inhibitors of CSF1R, such as PLX5622, selectively deplete microglia in vivo and in vitro by blocking trophic signaling mediated by CSF1 and IL-34(8,9). PLX5622 treatment therefore provides a robust and reproducible method for eliminating microglia from mixed glial cultures without disrupting astrocyte viability, cytoskeletal organization, or metabolic function.

In our optimized protocol, we integrated CSF1R inhibition into primary astrocyte workflows to markedly increase astrocyte purity and reduce confounding effects from microglia-derived factors. Our rationale was that, when combined with PDL-based adhesion, pipette-mediated mechanical dissociation, sequential OPC removal, and defined astrocyte culture conditions, this approach enables the generation of stable, high-fidelity astrocyte populations suitable for molecular, imaging-based, and biochemical assays. Importantly, the resulting cultures more accurately reflect astrocyte-intrinsic properties, providing an improved platform for studying astrocyte signaling pathways, inflammatory reactivity, trophic function, and receptor pharmacology.

A further advantage of this protocol is its compatibility with sex-specific analyses. Unlike many traditional astrocyte preparation methods, which require pooling tissue from multiple (often ≥4) neonatal brains to achieve adequate cell yield (10,11), our approach generates sufficient astrocyte numbers from a single mouse brain. This eliminates the need to pool sexes and enables researchers to derive independent male and female cultures without compromising cell yield or purity. Since sex influences glial development, inflammatory signaling, and susceptibility to neuropathology (12), the ability to generate high-quality astrocyte cultures from individual animals provides a critical methodological improvement for studies investigating sex-dependent astrocyte biology. Moreover, the ability to generate astrocyte cultures from a single mouse brain enables straightforward use of transgenic lines that require individual genotyping. This removes a major practical barrier associated with tissue pooling and greatly expands the range of genetically modified mouse models that can be studied using primary astrocyte cultures, including lines that were previously impractical to use in vitro due to the pooling strategy.

We present an optimized protocol for generating sex-defined, enriched primary astrocyte cultures from postnatal mouse cortices. By exposing the cultures to PLX5622 for 2 days to selectively deplete microglia, this method reliably produces highly purified astrocyte cultures with minimal contamination, enhancing consistency and reliability for downstream studies.

## 2. Methods

### 2.1 Materials

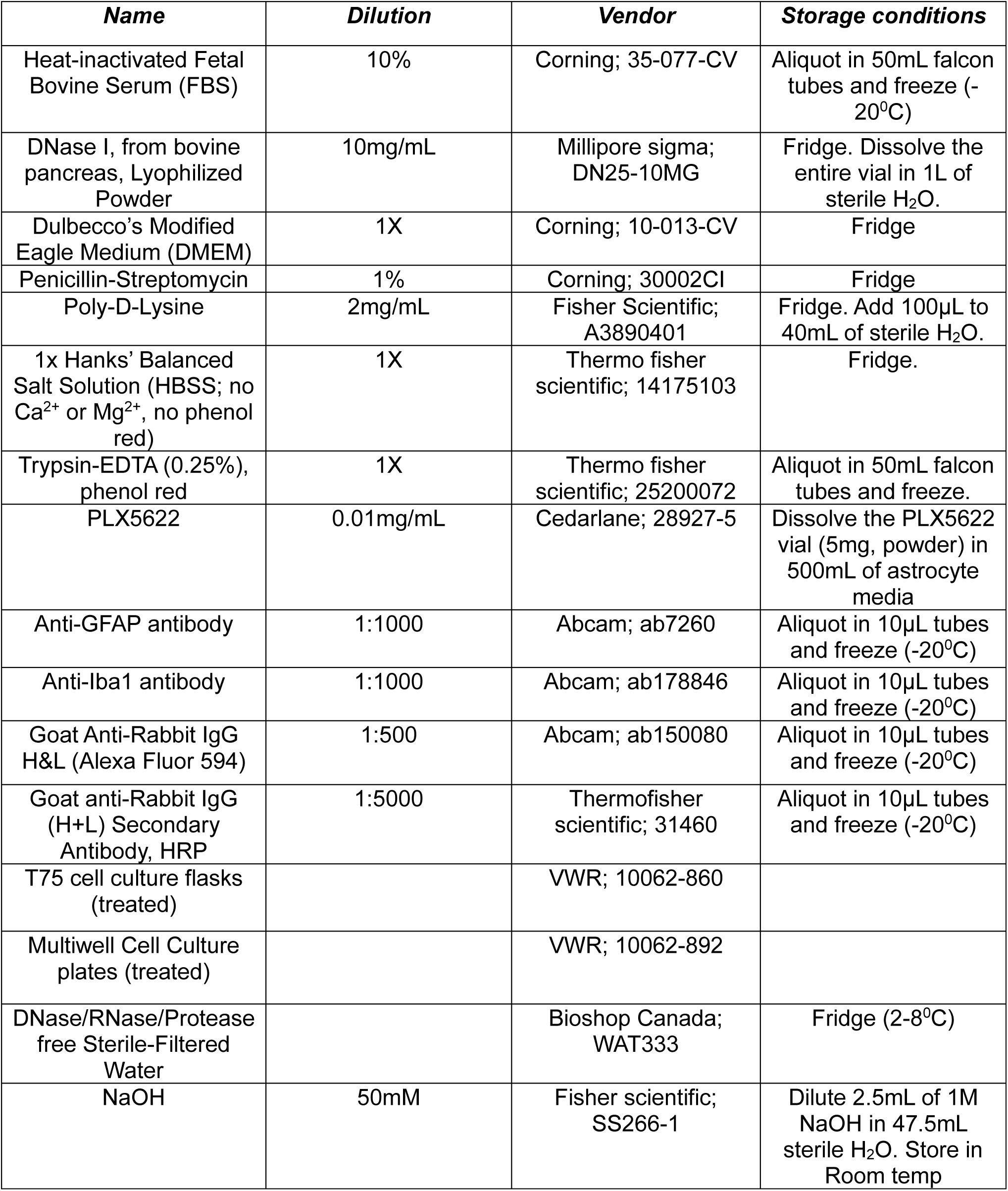
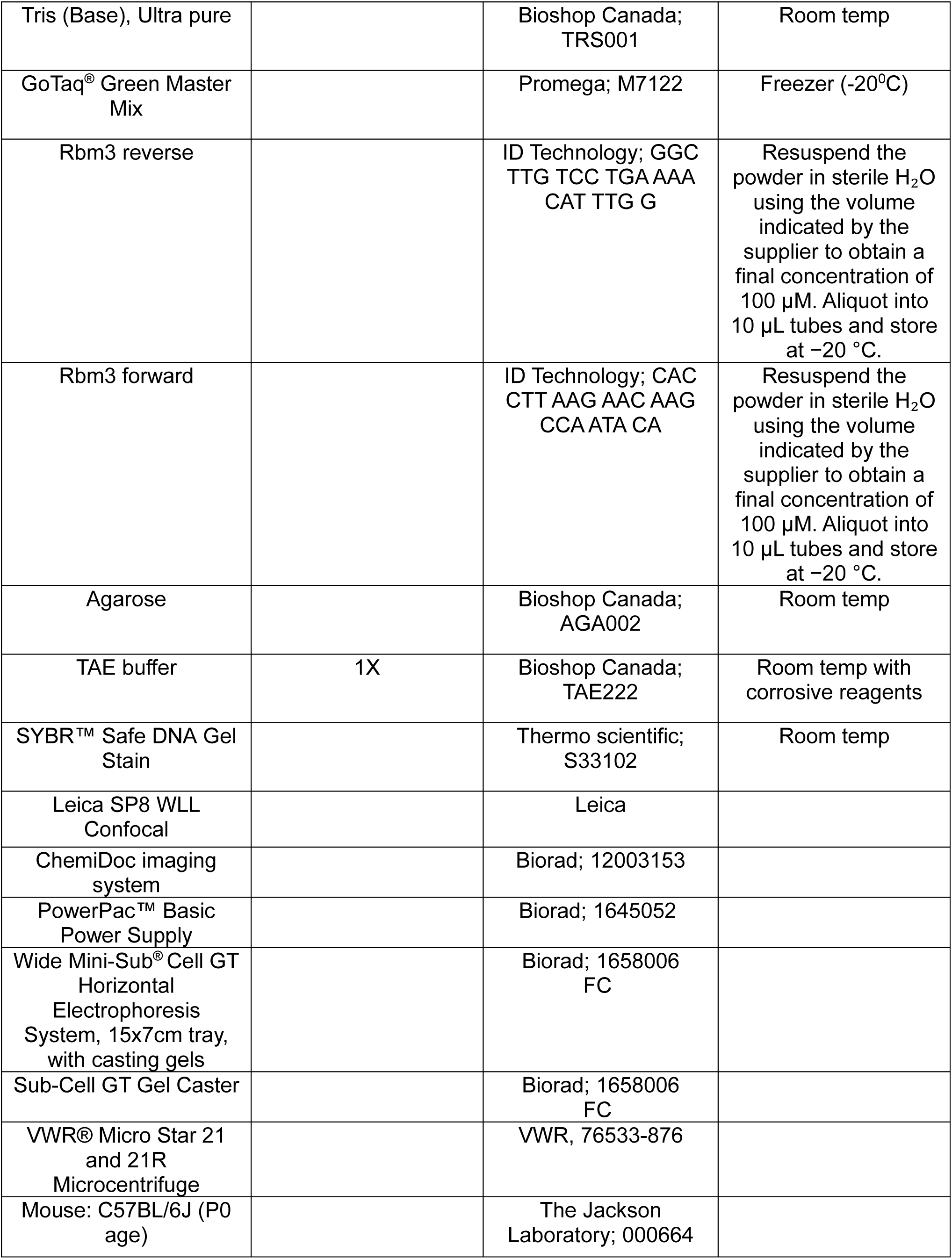
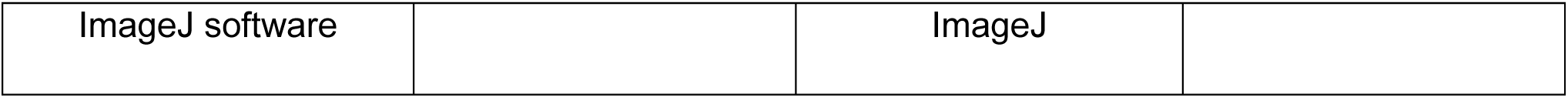

### 2.2 Procedure

#### 2.2.1 Coating Dishes with Poly-D-Lysine (PDL)

- Using the stock solution of 2 mg/mL, add 100 μL of PDL to 40 mL of water
- In treated T75 cell culture flasks, add 20mL PDL to cover the bottom of the flasks
- Incubate at 37°C for a minimum of 2 hours (ideally done in advance but T75 flasks can be prepared on the day off if required)
- After 2 hours, aspirate the PDL and wash the flasks with 4 mL of ddH_2_O (2x)
- Remove the water and let it dry with the cap off
  - **!CAUTION**: Coated flasks can be stored at 4°C for 1 month.

#### 2.2.2 Postnatal Astrocyte Media

To prepare astrocyte media

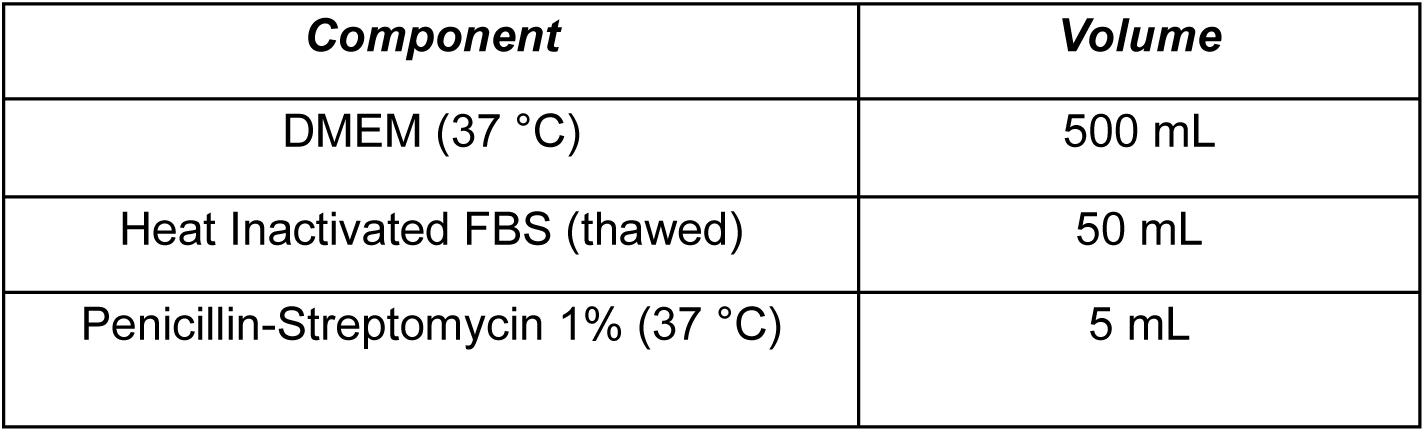

- **!CAUTION**: Astrocyte media should not be kept at 4°C for more than 1 month

#### 2.2.3 Dissecting embryonic brain

- Euthanize the P0-2 pup by decapitating with scissors (If material is needed for genotyping/ sex determination, cut off the tip of the pup’s tail post-decapitation and place in 1.5mL tubes).
- Place the head in a petri dish with cold HBSS.
- To remove the brain, first remove the skin from the top by placing the forceps in the middle and pulling towards the side
- With stabilizing forceps on the snout of the pup, use the other to remove the skull piece-by-piece.
  - **!CAUTION**: The skull is translucent and may be difficult to see and this step is preferably performed under dissecting microscope.
- Push the tip of one forceps underneath the skull and pull up.
- Break off small chunks and continue medially until cortices are reached
- To remove the brain, place the forceps underneath the cortices and lift out gently.
- Pinch off the olfactory bulbs and then the cerebellum and discard.
- Split the cortices down the longitudinal cerebral fissure (down the center).
- Remove the meninges by slipping the tip of the forceps underneath the membrane from the lateral side of the cortex (the meninges will contain all the vasculature. The brain is considered clean when there is no vasculature left)
- It may help to partially lift the brain out of the HBSS. The meninges will cling to the forceps, allowing for easier removal
- Place the brain in a 15 mL falcon tube containing 2 mL of astrocyte media

#### 2.2.4 Processing

- Using P1000 micropipette with a sterile tip, pipette up and down roughly 15 times, to break the brain down
- Once that is done, leave the falcon tube on ice for 5 minutes.
- Using another sterile P1000 tip, pipette again up and down for another 15-20 times to further break down the brain into pieces.
- Slowly remove the supernatant from the falcon tube and add it directly into the PDL-coated T75 flask. Add 8mL of astrocyte media into the flask and place in the incubator at 37 ^0^C 5% CO_2_
- Replace the astrocyte medium with fresh medium on day 1 and again on day 7.

#### 2.2.5 PLX5622 Astrocyte media preparation

- Prepare 500mL of fresh astrocyte culture media.
- Using the PLX5622 vial (5mg, powder), add approximately 1mL of the freshly prepared astrocyte medium to the vial. Gently invert the vial 2-3 times to ensure the powder is fully dissolved.
- Transfer the dissolved PLX5622 solution back into the 500mL of astrocyte medium.
  - **!CAUTION**: To ensure complete recovery of any remaining powder, add an additional 1mL of astrocyte medium to the PLX5622 vial and gently invert 2-3 times.
- Transfer the rinse back into the astrocyte medium.
- Gently invert the 500mL bottle of PLX5622-astrocyte medium 2-3 times to ensure thorough mixing and complete dissolution of PLX5622.

#### 2.2.6 Isolating enriched astrocytes (PLX5622 protocol)

- At 10-15 days the cells will be confluent and will contain a mixture of OPCs, microglia, and astrocytes
- Gently tap the sides of the flask 2-3 times to dislodge OPCs, which will detach from the surface and remain suspended in the supernatant.
- Aspirate the supernatant and gently wash cells with 10 mL of astrocyte media.
- Replace the media with another 10 mL containing 25 μL of DNAse I. Incubate at 37°C for 5 min
- Shake the flask horizontally by hand 20-30 times to remove the remaining OPCs
  - **!CAUTION**: It is important to shake vigorously enough to remove the OPCs but not the underlying astrocyte layer
- Remove the media and add 4 mL of warmed PBS to wash the cells twice.
- Add 12mL of PLX astrocyte media and allow to incubate for 24, 48, 72 or 96 hours (various exposure times were tested to identify the optimal duration, which was determined to be 2 days).
- After the designated incubation period, aspirate PLX astrocyte media and replace it with regular astrocyte media.
- If the flask is 90-95% confluent, split the flasks into desired dishes.

#### 2.2.7 Maintaining and Feeding Astrocyte Cultures

- Every 4 days, replace the media with fresh astrocyte media (12mL).

### 2.3 Sex determination

- The protocol was adapted from (13,14), with the modifications noted below.

#### 2.3.1 Extracting DNA from tails

- Add 250µL of 50mM NaOH to each tail in 1.5mL tubes. Incubate the tissue on a heat block at 95°C for 45mins
- After 45mins, let the tubes cool for a few minutes, until roughly 50-70°C. Vortex tubes to digest the tissue, ensuring that they contain only hair and some cartilage remnants, as soft tissue should have dissolved completely
- Add 250µL of 0.5M Tris-HCl (pH 5.5) to neutralize the samples. Vortex to mix
  - Tris-HCl is made by dissolving approximately 10.1g of Tris in 100mL of sterile H_2_O, then adding HCl (dropwise) to reach a final pH of 5.5
  - Centrifuge the tubes at 14,800 RPM for 2mins.

#### 2.3.2 PCR preparation (sex-typing)

| Reagent | Volume (µL) |
| --- | --- |
| Sterile H<sub>2</sub>O | 10 |
| GoTaq® Green Master Mix | 12.5 |
| 100uM Rbm3 reverse | 1.5 |
| 100uM Rbm3 forward | 1.5 |
| Tail DNA | 3 |

- To prepare the primers, resuspend them in sterile H_2_O as per the indicated volumes by the supplier to achieve 100µM of final concentration.
- Add the indicated volumes to PCR tubes
- Use the following PCR protocol for DNA amplification:
  A. 94°C for 3mins
  B. 94°C for 30 seconds
  C. 65°C for 30 seconds
  D. 68°C for 45 seconds
    - **10 cycles of steps A-D**
  E. 94°C for 30 seconds
  F. 60°C for 30 seconds
  G. 72°C for 45 seconds
    - **20 cycles of steps E-G**
  H. 72°C for 5mins
  I. Hold at 4°C

#### 2.3.3 Gel electrophoresis

- Prepare agarose gel by dissolving 3g agarose in 100 mL sterile water supplemented with 2 mL 50X TAE buffer. Heat the solution in 30-second microwave intervals until fully dissolved and clear.
- Pour into gel electrophoresis cassette, add comb, and leave to solidify for approximately 20mins
- Transfer into gel electrophoresis tank and add running buffer to cover
  - To make 1X running buffer, add 20mL of 50X TAE buffer in 800mL of ddH_2_O
- Run the gel at 90 volts for 61mins.
- Image the gel using Chemidoc using the nucleic acid Sybrsafe module.

### 3. Expected results

PLX5622 is a selective inhibitor of CSF1R widely used to achieve efficient and reversible microglial depletion in the CNS. By blocking CSF1R signaling, PLX5622 enables precise experimental investigation of microglial contributions to neural homeostasis and disease (15–18). We propose a novel application of PLX5622, used in combination with established methodologies, to generate highly pure, enriched primary astrocyte cultures with defined sex specificity. To optimize this process, we experimented with varying days of PLX5622 treatment followed by immunocytochemistry to ascertain what treatment duration gives rise to total microglia depletion while minimizing any potential off target effects or toxicity. Conditions ranged from 1 day to 4 days of treatment with 0.01mg/mL PLX5622 in astrocyte media with each culture being wholly comprised of cells of a specific sex. Efficacy of the enrichment process was evaluated with astrocyte and microglia markers, GFAP and Iba1, respectively, in immunostaining experiments whilst PCR and gel electrophoresis is used to determine sex of origin animal and derived culture. The respective purity of cultures and presence of astrocytes and microglia is visualised via immunofluorescence imaging (**Fig 1, 2, 3, 4**).

**Figure 1.**
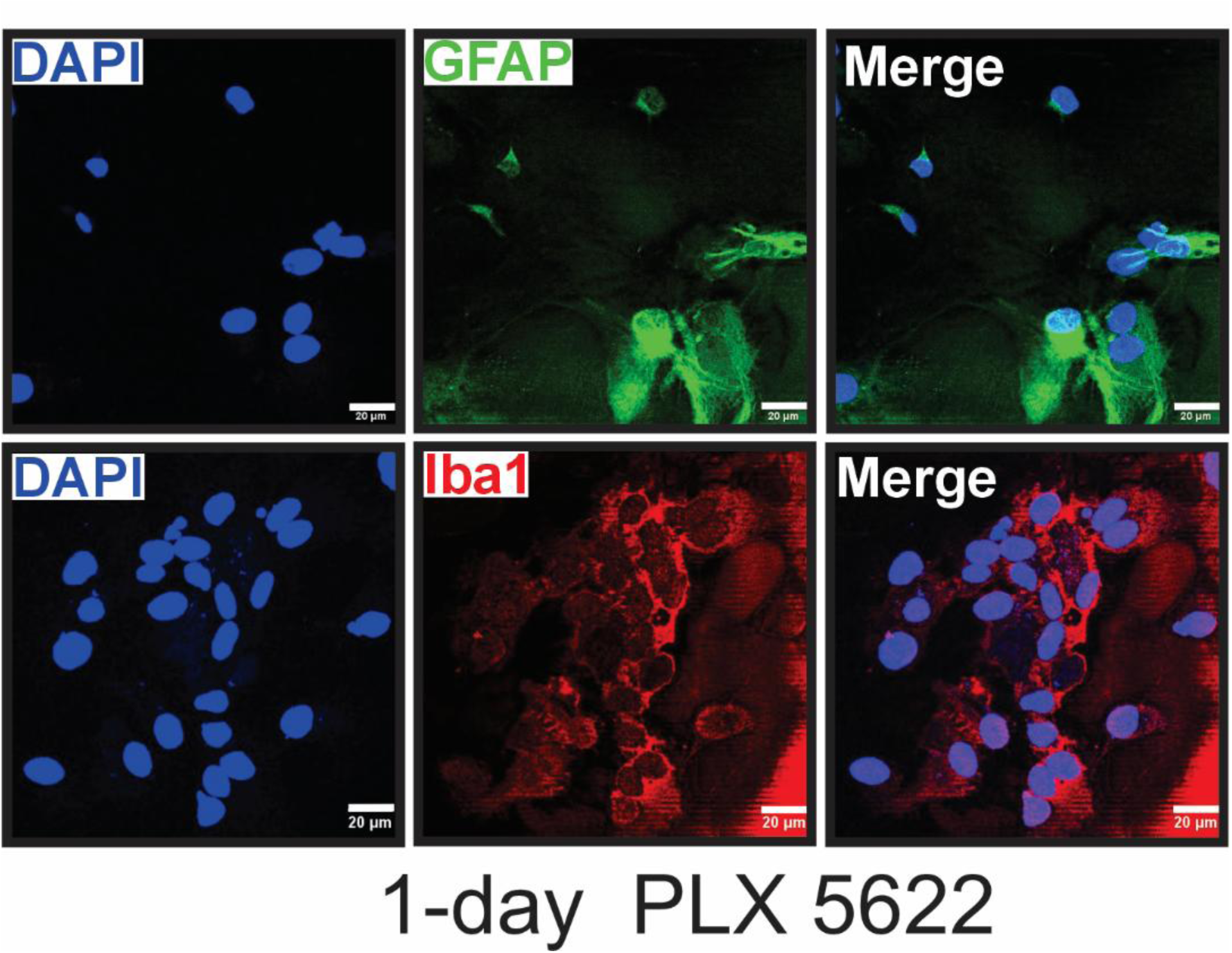
Astrocyte and microglia immunofluorescence images in PLX5622 1-day treatment condition. Representative immunofluorescence images of astrocytes and microglia in primary astrocyte cultures in 1-day PLX5622 (0.01mg/ml) treatment condition. With 24 hours of PLX5622 treatment, images depict presence of both astrocytes and microglia in culture and thus indicate incomplete depletion of microglia. Nuclei of all cells were stained with 4’,6-diamidino-2-phenylindole (DAPI) and shown in red. GFAP-positive astrocytes were identified with GFAP markers and shown in green, microglia identified with Iba-1 markers and shown in green. Immunofluorescence images were acquired with a 63x objective in Leica confocal microscope (SP8 WLL). All images processed in ImageJ software.

Immunocytochemical examination of the cultures indicated that 24 hours of PLX5622 treatment was not sufficient to fully eliminate the presence of microglia from cultures, whilst there is a marked decrease in microglia presence, contamination is nevertheless still present in the culture (stained using Iba1 marker; **Fig 1**). At the two-day mark of PLX5622 treatment, we can observe total depletion of microglia in culture with astrocyte count and morphology unaffected (**Fig. 2**). In the three-day and four-day conditions, we similarly observe total depletion of microglia, however, astrocyte count does also appear to show a decline correlated with treatment duration (**Fig. 3 and 4**). This may suggest off-target effects or some degree of cytotoxicity and cell death driven by PLX5622 or a downstream effect. As such, the two-day condition is not only optimal for efficiency but also gives the highest yield of pure astrocytes. These findings are congruent with immunoblots (**Fig. 5**), also displaying the presence of Iba1 in the absence of PLX5622 and lack thereof in PLX5622-treated conditions. Together, PLX5622 treatment induces a time-dependent depletion of microglia in primary astrocyte cultures, as confirmed by immunofluorescence and immunoblot analyses. ***An exposure duration of 2 days was optimal for achieving highly pure astrocyte cultures while minimizing cytotoxicity***.

**Figure 2.**
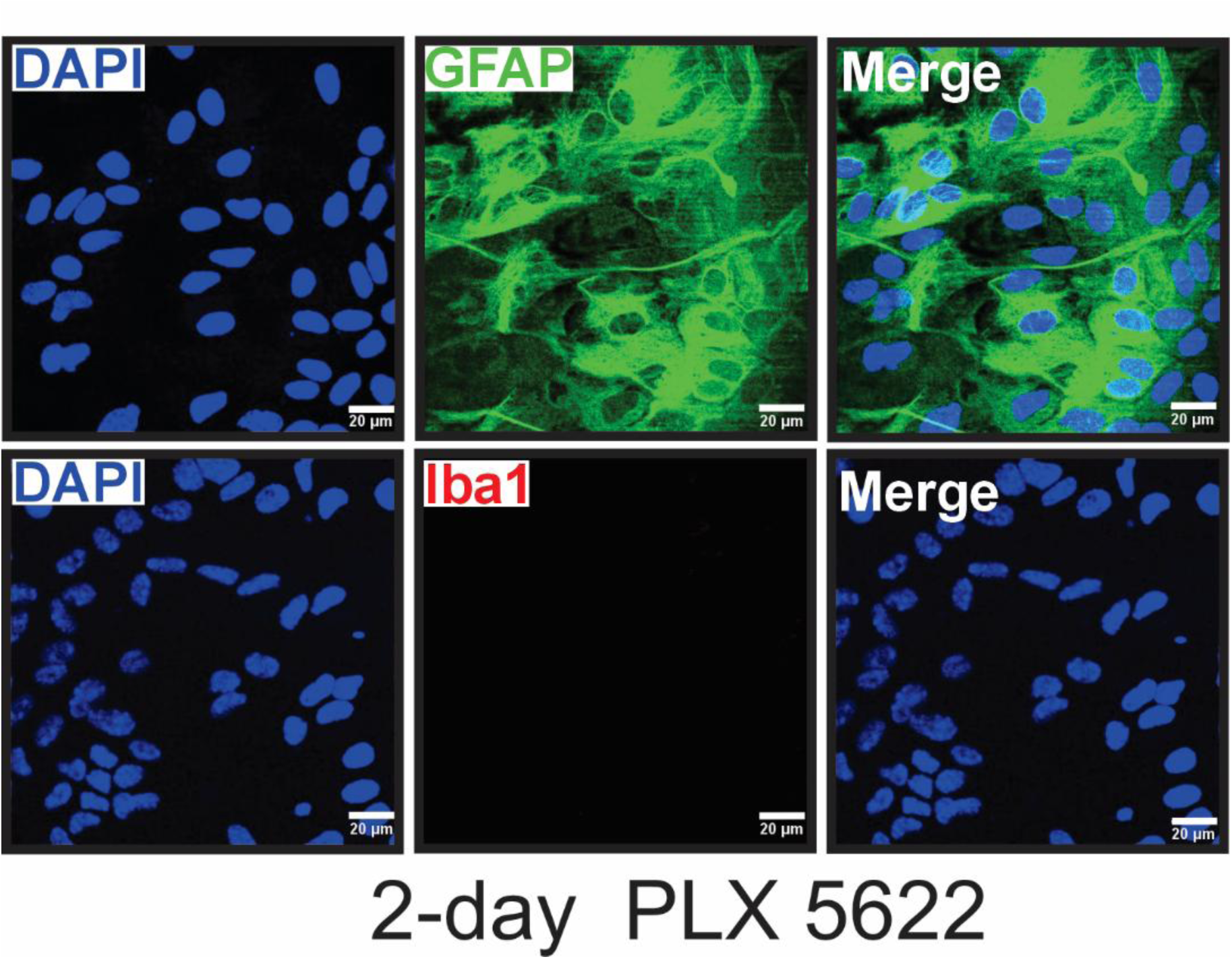
Astrocyte and microglia immunofluorescence images in PLX5622 2-day treatment condition. Representative immunofluorescence images of astrocytes and microglia in primary astrocyte cultures in 2-day PLX5622 (0.01mg/ml) treatment condition. Images derived from cultures treated with 48 hours of PLX5622 appear to result in total depletion of microglia in culture with astrocyte numbers and morphology completely unaffected, strongly supporting 2 days as the optimal treatment duration. Nuclei of all cells were stained with 4’,6-diamidino-2-phenylindole (DAPI) and shown in red. GFAP-positive astrocytes were identified with GFAP markers and shown in green, microglia identified with Iba-1 markers and shown in green. Fluorescent images were acquired with a 63x objective in Leica confocal microscope (SP8 WLL). All images processed in ImageJ software.

**Figure 3.**
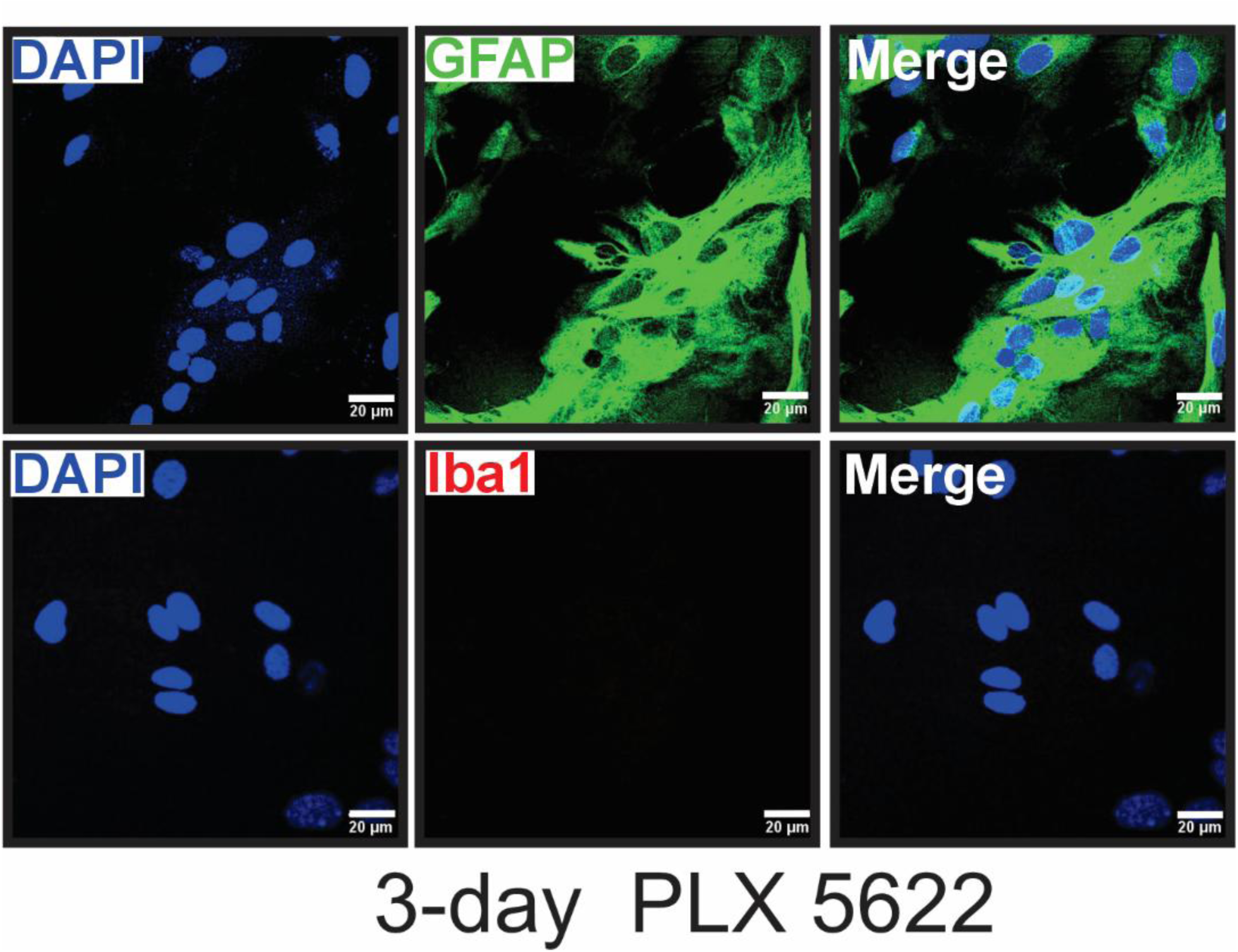
Astrocyte and microglia immunofluorescence images in PLX5622 3-day treatment condition. Representative immunofluorescence images of astrocytes and microglia in primary astrocyte cultures in 3-day PLX5622 (0.01mg/ml) treatment condition. In the 72-hour PLX5622 treatment condition, we can observe a thorough depletion of microglia with astrocytes relatively unaffected, not dissimilar to the 2-day condition. Nuclei of all cells were stained with 4’,6-diamidino-2-phenylindole (DAPI) and shown in red. GFAP-positive astrocytes were identified with GFAP markers and shown in green, microglia identified with Iba-1 markers and shown in green. Fluorescent images were acquired with a 63x objective in Leica confocal microscope (SP8 WLL). All images processed in ImageJ software.

**Figure 4.**
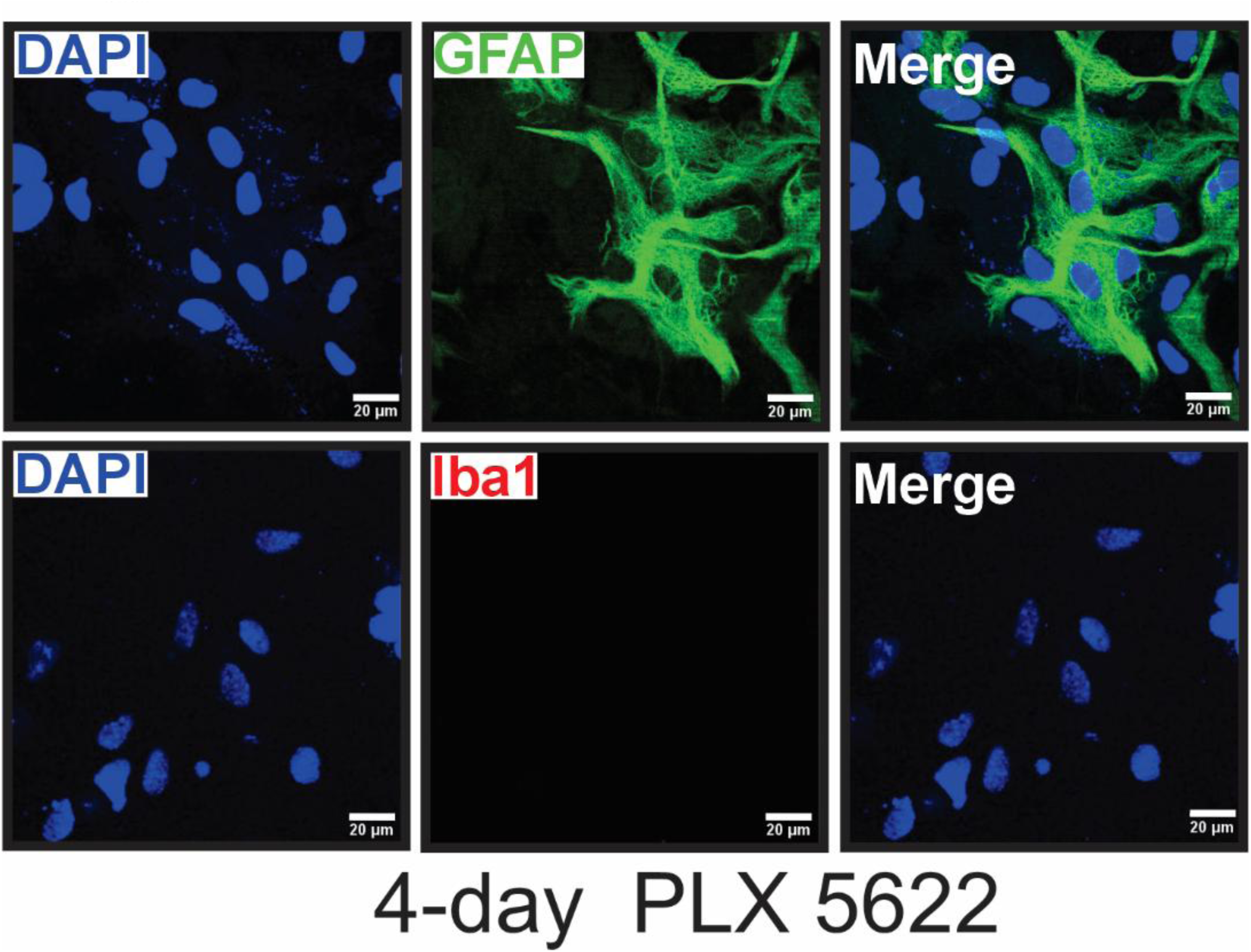
Astrocyte and microglia immunofluorescence images in PLX5622 4-day treatment condition. Representative immunofluorescence images of astrocytes and microglia in primary astrocyte cultures in 4-day PLX5622 (0.01mg/ml) treatment condition. Consistent with other treatment conditions that indicate the time-dependent action of PLX5622, the 96-hour treatment condition also results in removal of microglial presence in culture, leaving a purified astrocyte culture. However, in the 4-day condition, there is discernable impact to total cell count, suggesting some degree of potential cytotoxic effect from prolonged exposure. Nuclei of all cells were stained with 4’,6-diamidino-2-phenylindole (DAPI) and shown in red. GFAP-positive astrocytes were identified with GFAP markers and shown in green, microglia identified with Iba-1 markers and shown in green. Fluorescent images were acquired with a 63x objective in Leica confocal microscope (SP8 WLL). All images processed in ImageJ software.

**Figure 5.**
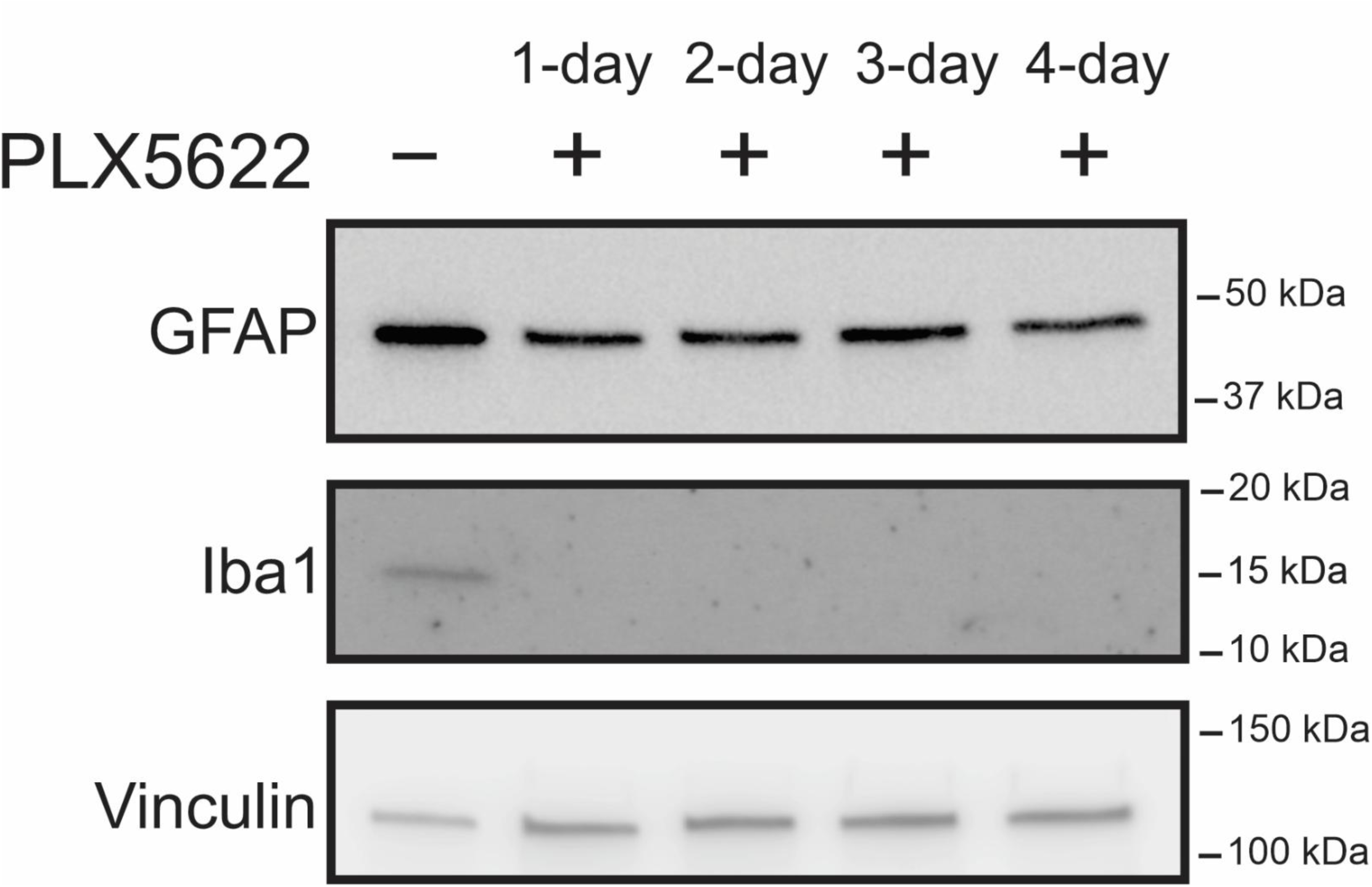
GFAP and Iba1 immunoblots under various PLX5622 treatment conditions. Representative immunoblots showing the protein expression of GFAP (using anti-GFAP; 1:1000) and Iba1 (using anti-Iba1; 1:1000) in the absence and the presence of various PLX5622 treatments. In PLX5622-treated samples, Iba1 positive cells are absent at days 1, 2, 3, and 4, indicating its efficacy in depleting microglia compared with untreated controls. In contrast, GFAP positive astrocytes are consistently present across all conditions, regardless of PLX5622 exposure. This demonstrates selective depletion of microglia without significant loss of astrocytes.

Alongside immunoblotting and immunocytochemical analysis, in a process known as sex-typing, a simplex PCR using primers for the X-linked Rbm31x gene and Y-linked gametolog Rbm31y is conducted on genetic material (tails) derived from the source animals that are respectively used for sex-specific astrocyte cultures (14). This process of sex-typing is a crucial part of the protocol; while the usage of PLX5622 for consistent microglial depletion has been highlighted repeatedly, the strength of the protocol also lies in its ability to produce cultures of cells comprised entirely of one sex. As such, it is imperative that genotyping (sex-typing) is performed to identify and capitalise on the protocol’s ability to give rise to sex-specific in-vitro research models. The DNA amplification process by the PCR is followed by gel electrophoresis and imaging for sex determination to identify the specific sex of isolated and purified cell cultures (**Fig. 6**). We observe an expected and consistent result whereby each animal and thus, each culture, gives rise to clear, clean bands, with 2 bands indicating a male animal and subsequent culture and 1 band indicating a female animal and subsequent culture.

**Figure 6.**
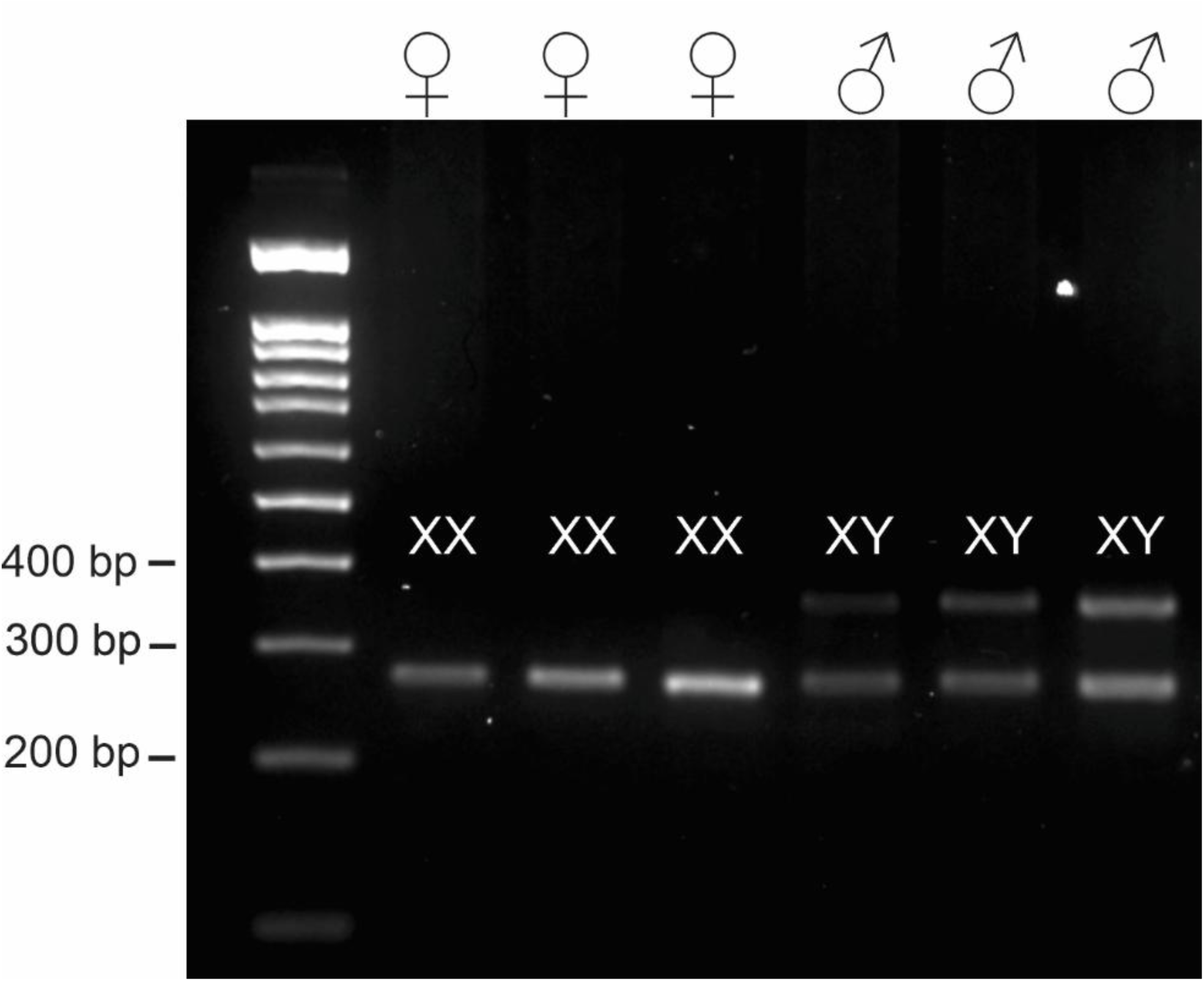
PCR Gel electrophoresis for sex determination. PCR Gel Electrophoresis performed on tails derived from the same neonates used to produce primary astrocyte cultures. Rbm31 primers are used to target the Rbm31x and Rbm31y genes on the X and Y chromosomes. One band indicates female (XX) and two bands indicates male (XY).

### 4. Troubleshooting

#### 4.1 Incomplete Microglial Depletion Following PLX5622 Treatment

##### Possible causes

Incomplete dissolution of PLX5622, insufficient exposure time, or high initial microglial density may result in residual microglia following treatment.

##### Solutions

Ensure PLX5622 is fully dissolved prior to addition to culture media, with no visible particulates. Confirm the final working concentration is 0.01 mg/mL and maintain treatment for a full 48 hours. Avoid overconfluent cultures prior to treatment, as increased cell density may reduce drug efficacy.

#### 4.2 Astrocyte loss or reduced viability after PLX5622 treatment

##### Possible causes

Extended PLX5622 exposure, incomplete media replacement following treatment, or excessive mechanical stress during handling may reduce astrocyte viability.

##### Solutions

Limit PLX5622 exposure strictly to 48 hours. Replace PLX5622-astrocyte medium promptly with fresh astrocyte medium after treatment. Minimize time outside the incubator and avoid excessive washing or agitation during media changes.

#### 4.3 Variable astrocyte yield between preparations

##### Possible causes

Inconsistent dissection technique, incomplete removal of meninges, or variability of mechanical dissociation can affect final cell yield.

##### Solutions

Ensure complete removal of meninges, which contain non-glial contaminants. Standardize trituration force and number of pipetting steps. Use only P0 pups and maintain consistency in dissection timing and handling.

#### 4.4 Poor cell attachment or uneven astrocyte monolayers

##### Possible causes

Inadequate PDL coating, improper storage of coating flasks, or insufficient drying prior to use.

##### Solutions

Prepare fresh PDL solution and ensure complete coverage of culture surfaces. Allow coated flasks to dry fully before use and store at 4^0^C for no longer than one month. Avoid disturbing flasks during the initial attachment period.

### 5. Discussion

This protocol presents a consistent and efficient method at producing a pure primary sex-specific astrocyte culture derived from P0-2 mouse cortices. Using neonatal (P0-2) cortices from wildtype C57BL/6J mice, we can derive a mixed primary culture of cells that can be purified to produce an enriched sex-specific primary astrocyte culture for in-vitro study. It is a multi-step process that specifically targets two main glial cell populations that are often pervasive in primary cell cultures, oligodendrocyte precursor cells and microglia. The primary step of decapitation and brain dissection follows the established protocol(10), which has proven efficacy and yields consistent results in producing a mixed primary glial culture via adhesion on PDL-coated substrates.

OPCs are first selectively removed from the mixed primary cell culture through capitalization of differential adhesion properties between the glial cell types(19). As OPCs are not as adherent as the microglia and astrocytes in culture, they can be detached from the flask surface via shaking and tapping. Detached OPCs float in the supernatant and can subsequently be aspirated and removed from the mixed primary culture. Conversely, this also serves as an effective method to obtain a purified OPC primary culture if the aspirated supernatant is subsequently plated and cultured.

Following removal of OPCs, microglia need to also be eliminated to leave a purified astrocyte culture. Traditional methods struggle with microglia removal and for the most part is also reliant on differential adhesion properties and multistage shaking to suspend desired cell types(10,11). This not only proves to be a strenuous process, but it also risks loss of astrocytes or incomplete depletion of microglia. The aforementioned methods are based on previously established protocols which underlies the foundation of isolating primary cells from neonatal mouse pups. However, this protocol notably incorporates the usage of CSF1R inhibitor, PLX5622, to refine and accelerate the process of microglia depletion in mixed primary glial cell cultures. In current prevailing protocols, the final purification step for astrocytes is still reliant on differential adhesion properties between microglia and astrocytes and involves trypsin-ising astrocytes to suspend and isolate them. While effective, this process may run the risk of inadvertent cell death(20) and can limit clear separation of microglia and astrocytes, given their relatively similar adhesion characteristics on PDL-coated surfaces. Critically, our utilization of PLX5622 greatly minimises the risk of astrocytic cell death and yields pure primary astrocytes in a highly reproducible manner.

Additionally, traditional protocols follow a similar workflow that requires multiple (in some cases up to four) mouse neonatal cortices to achieve appropriate astrocyte density in tissue culture (10,11). In contrast, pivotally, our process of microglial depletion with PLX5622 increases the yield of astrocytes such that one mouse pup can afford equivalent astrocyte density for a T75 tissue culture flask. This is expectedly beneficial for increased efficiency and reduced animal use but more meaningfully presents the added benefit of enabling more robust in-vitro research into sex- and genotype-specific differences. As purified astrocyte cultures are not comprised of a mixture of cells from four animals but is rather a 1:1 representation of one animal, the cell culture derived is in turn wholly composed of and representative of the sex and genotype of this animal. As such, any results from subsequent work on the cells would allow for more robust in-vitro research into sex and genotype-specific phenomena.

The cornerstone of this protocol, PLX5622, is not without substitutes, namely PLX3397, another CSF1R inhibitor. PLX3397 is another readily available CSF1R inhibitor that excels in microglia depletion in vitro and in vivo. As CSF1R inhibitors display variable specificity, a direct comparison was made showing that PLX5622 eclipses PLX3397 both in target specificity and in efficacy of microglial depletion(21). More specifically, PLX3397 exhibited consistently lower percentages of microglia depletion at all time points and concentrations whilst also causing off-target effects of cytotoxicity, significantly reducing levels of OPCs even at low concentrations. Even though OPCs reduction is a desirable outcome in our primary astrocyte purification protocol, the uncontrolled and incomplete nature of its action (only 30-50% OPC depletion) coupled with the risk of cytotoxicity make it a less suitable CSF1R inhibitor for our protocol. In contrast, with its greater selectivity and safer, thorough microglial depletion, we deemed PLX5622 to be more fitting for this protocol. However, it must be noted that these direct comparisons were all undertaken either in vivo or ex vivo, there has yet to be any direct in vitro comparisons of the two CSF1R inhibitors.

While in vitro astrocyte models are invaluable for investigating CNS physiology and disease mechanisms, this protocol has inherent limitations. First, like other primary cell culture methods, it is time-sensitive and relies on postnatal day 0-2 mouse pups, restricting flexibility and scalability. Second, despite documented differences in adhesion properties between astrocytes and OPCs, there remains a small but unavoidable risk of astrocyte loss during the shaking process, which is difficult to control or quantify.

In summary, in today’s culture of ever-growing interest in astrocytes, this protocol serves as a powerful tool in garnering a better understanding of astrocytic mechanisms and biology. The diverse research applicability of a pure, sex-specific astrocyte culture in vitro cannot be overstated and will undoubtedly be used as the foundation for numerous advancements in research. However, it is important to note that these purified cultures are not necessarily representative of in-vivo conditions. Consequently, we must carefully consider the transferability of findings and not forgo other methods to use in tandem for more robust, holistic research.

### Declarations

#### Ethics approval and consent to participate

All procedures were carried out in the University of British Columbia Preclinical Discovery Centre in accordance with the Canadian Council on Animal Care and approved by the University of British Columbia Committee on Animal Care (protocols, A23-0274, A23-0045).

#### Availability of data and materials

The datasets used and/or analysed during the current study are available from the corresponding author on reasonable request.

#### Author contributions

S.A. analyzed and interpreted the data, contributed to conceptualization and visualization, and writing of the manuscript. E.A. contributed to conceptualization, data analysis, visualization, and writing of the manuscript. J.A.W. performed data analyses and visualization. K.S.A-E supervised the project, contributed to conceptualization and manuscript review and editing, and secured funding. All authors read and approved the final manuscript.

## Acknowledgements

K.S.A-E is a Michael Smith Health Research BC funded Health-Professional Investigator. Graphical abstract was created using BioRender.com.

## Abbreviations

CNS: Central Nervous System
CSF1R: colony-stimulating factor 1 receptor
DMEM: Dulbecco’s Modified Eagle Medium
FBS: Heat-inactivated Fetal Bovine Serum
HBSS: Hanks’ Balanced Salt Solution
OPC: Oligodendrocyte Progenitor Cells
PDL: Poly-D-Lysine

## References

1. Siracusa R, Fusco R, Cuzzocrea S. Astrocytes: Role and Functions in Brain Pathologies. Front Pharmacol. 2019;10:1114.

2. Sofroniew M V. Astrocyte barriers to neurotoxic inflammation. Nat Rev Neurosci. 2015 May 20;16(5):249–63.

3. Ibrahim AM, Pottoo FH, Dahiya ES, Khan FA, Kumar JBS. Neuron-glia interactions: Molecular basis of alzheimer’s disease and applications of neuroproteomics. Eur J Neurosci. 2020 Jul;52(2):2931–43.

4. De Pittà M, Brunel N, Volterra A. Astrocytes: Orchestrating synaptic plasticity? Neuroscience. 2016 May 26;323:43–61.

5. Liddelow SA, Guttenplan KA, Clarke LE, Bennett FC, Bohlen CJ, Schirmer L, et al. Neurotoxic reactive astrocytes are induced by activated microglia. Nature. 2017 Jan 26;541(7638):481–7.

6. Nutma E, van Gent D, Amor S, Peferoen LAN. Astrocyte and Oligodendrocyte Cross-Talk in the Central Nervous System. Cells. 2020 Mar 3;9(3).

7. Saura J. Microglial cells in astroglial cultures: a cautionary note. J Neuroinflammation. 2007 Oct 15;4:26.

8. Spangenberg E, Severson PL, Hohsfield LA, Crapser J, Zhang J, Burton EA, et al. Sustained microglial depletion with CSF1R inhibitor impairs parenchymal plaque development in an Alzheimer’s disease model. Nat Commun. 2019 Aug 21;10(1):3758.

9. Mein N, von Stackelberg N, Wickel J, Geis C, Chung HY. Low-dose PLX5622 treatment prevents neuroinflammatory and neurocognitive sequelae after sepsis. J Neuroinflammation. 2023 Dec 1;20(1):289.

10. Güler BE, Krzysko J, Wolfrum U. Isolation and culturing of primary mouse astrocytes for the analysis of focal adhesion dynamics. STAR Protoc. 2021 Dec 17;2(4):100954.

11. Schildge S, Bohrer C, Beck K, Schachtrup C. Isolation and culture of mouse cortical astrocytes. J Vis Exp. 2013 Jan 19;(71).

12. Müller L, Di Benedetto S, Müller V. Influence of biological sex on neuroinflammatory dynamics in the aging brain. Front Aging Neurosci. 2025 Aug 29;17.

13. Tunster SJ. Genetic sex determination of mice by simplex PCR. Biol Sex Differ. 2017 Oct 17;8(1):31.

14. Abd-Elrahman KS, Albaker A, de Souza JM, Ribeiro FM, Schlossmacher MG, Tiberi M, et al. Aβ oligomers induce pathophysiological mGluR5 signaling in Alzheimer’s disease model mice in a sex-selective manner. Sci Signal. 2020 Dec 15;13(662).

15. Henry RJ, Ritzel RM, Barrett JP, Doran SJ, Jiao Y, Leach JB, et al. Microglial Depletion with CSF1R Inhibitor During Chronic Phase of Experimental Traumatic Brain Injury Reduces Neurodegeneration and Neurological Deficits. The Journal of Neuroscience. 2020 Apr 1;40(14):2960–74.

16. He Y, Liu T, He Q, Ke W, Li X, Du J, et al. Microglia facilitate and stabilize the response to general anesthesia via modulating the neuronal network in a brain region-specific manner. Elife. 2023 Dec 22;12.

17. Császár E, Lénárt N, Cserép C, Környei Z, Fekete R, Pósfai B, et al. Microglia modulate blood flow, neurovascular coupling, and hypoperfusion via purinergic actions. Journal of Experimental Medicine. 2022 Mar 7;219(3).

18. Profaci CP, Harvey SS, Bajc K, Zhang TZ, Jeffrey DA, Zhang AZ, et al. Microglia are not necessary for maintenance of blood-brain barrier properties in health, but PLX5622 alters brain endothelial cholesterol metabolism. Neuron. 2024 Sep;112(17):2910–2921.e7.

19. O’Meara RW, Ryan SD, Colognato H, Kothary R. Derivation of enriched oligodendrocyte cultures and oligodendrocyte/neuron myelinating co-cultures from post-natal murine tissues. J Vis Exp. 2011 Aug 21;(54).

20. Lordon B, Campion T, Gibot L, Gallot G. Impact of trypsin on cell cytoplasm during detachment of cells studied by terahertz sensing. Biophys J. 2024 Aug;123(16):2476–83.

21. Liu Y, Given KS, Dickson EL, Owens GP, Macklin WB, Bennett JL. Concentration-dependent effects of CSF1R inhibitors on oligodendrocyte progenitor cells ex vivo and in vivo. Exp Neurol. 2019 Aug;318:32–41.

